# Injectable hydrogels for sustained co-delivery of subunit vaccines enhance humoral immunity

**DOI:** 10.1101/2020.05.26.117465

**Authors:** Gillie A. Roth, Emily C. Gale, Marcela Alcántara-Hernández, Wei Luo, Eneko Axpe, Rohit Verma, Qian Yin, Anthony C. Yu, Hector Lopez Hernandez, Caitlin L. Maikawa, Anton A. A. Smith, Mark M. Davis, Bali Pulendran, Juliana Idoyaga, Eric A. Appel

**Author notes:** Person to whom correspondence should be addressed. **Author Contributions**: G.A.R., E.C.G., M.A.H., W.L., E.A., R.V., Q.Y., M.M.D, B.P., J.I., and E.A.A designed experiments; G.A.R., E.C.G, M.A.H., W.L., E.A., R.V., A.C.Y., C.L.M., and A.A.A.S. conducted experiments; G.A.R., E.C.G., M.A.H., W.L., E.A., R.V., H.L.H., and E.A.A. analyzed data; and G.A.R. and E.A.A. wrote paper.

## Abstract

Vaccines aim to elicit a robust, yet targeted, immune response. Failure of a vaccine to elicit such a response arises in part from inappropriate temporal control over antigen and adjuvant presentation to the immune system. In this work, we sought to exploit the immune system’s natural response to extended pathogen exposure during infection by designing an easily administered slow-delivery vaccine platform. We utilized an injectable and self-healing polymer-nanoparticle (PNP) hydrogel platform to prolong the co-delivery of vaccine components to the immune system. We demonstrated that these hydrogels exhibit unique delivery characteristics whereby physicochemically distinct compounds (such as antigen and adjuvant) could be co-delivered over the course of weeks. When administered in mice, hydrogel-based sustained vaccine exposure enhanced the magnitude, duration, and quality of the humoral immune response compared to standard PBS bolus administration of the same model vaccine. We report that the creation of a local inflammatory niche within the hydrogel, coupled with sustained release of vaccine cargo, enhanced the magnitude and duration of germinal center responses in the lymph nodes. This strengthened germinal center response promoted greater antibody affinity maturation, resulting in a more than 1000-fold increase in antigen-specific antibody affinity in comparison to bolus immunization. In summary, this work introduces a simple and effective vaccine delivery platform that increases the potency and durability of subunit vaccines.

## Main Article

Vaccines are critical for combating infectious diseases across the globe. Unfortunately, vaccines against many infectious diseases have proven to be difficult to develop for various reasons ranging from poor antigen immunogenicity, poor breadth of protection against rapidly mutating antigenic targets, or poor durability of immune protection^1-4^. To elicit a durable, protective antibody response, vaccines must interact with the appropriate cell types at the right time and place while also providing the necessary cues to guide the immune system^5^. With regards to temporal dynamics, natural infections expose the immune system to antigen and inflammatory signals for 1-2 weeks^5^. Conversely, the short-term presentation of subunit vaccines from a single bolus administration persists for only 1-2 days^5^. The sustained release of soluble antigen provides essential signals to germinal centers (GCs) in the lymph nodes—the structures responsible for the affinity maturation of B cells—leading to high affinity antibody production and generation of humoral immune memory^6-8^. Recent work demonstrates that slow delivery of an HIV vaccine using surgically implantable osmotic pumps in nonhuman primates resulted in higher neutralizing titers^9^. Similarly, sustained release of a stabilized HIV antigen from a microneedle patch led to increased serum titers and bone marrow plasma cells in mice^10,11^. It has been reported that the mechanism of traditional adjuvant systems such as aluminum hydroxide (Alum) rely on increased antigen persistence in the body^12-14^, a feature that was recently exploited further through engineering antigens that adhere more strongly to Alum particles, thereby providing sustained antigen delivery to lymph nodes and improving the humoral response^15^. These studies and others demonstrate that the kinetics of antigen presentation to the immune system dramatically influence the adaptive immune response^9,10,12,13,16-27^. However, many previously reported sustained-release technologies are limited by several factors: (i) the inability to deliver chemically diverse cargo, (ii) restricted time-frame tunability for prolonged release, and/or (iii) low yielding and, oftentimes, challenging multi-step syntheses^28^. Therefore, there is a significant need for the development of new controlled delivery approaches that allow for the sustained exposure of vaccine components to the immune system.

Hydrogel materials are well-suited for a variety of biomedical applications due to their high water content and mechanical tunability that make them similar to many biological tissues^29,30^. They have been used as cell culture substrates^31-33^, tissue engineering or tissue regeneration templates^34,35^, and drug delivery vehicles^36,37^. More recently, hydrogels for controlled delivery of both cancer and infectious disease immunotherapies have shown promising results^38,39^. Numerous approaches have been utilized to create hydrogel systems for vaccine delivery^40-43^, though often focusing on innate immune activation for cancer therapy rather than sustained vaccine delivery to enhance humoral immune responses.

Importantly, traditional covalently cross-linked hydrogels are often not appropriate for vaccine applications as they cannot be easily administered and have limitations in their delivery kinetics for diverse cargo^29^. Polymer-nanoparticle (PNP) hydrogels are a type of supramolecular hydrogel where the polymeric constituents are held together by dynamic, multivalent non-covalent interactions between polymers and nanoparticles^44-47^. These materials exhibit many favorable characteristics such as high drug loading capacity, gentle conditions for encapsulation of biologic cargo, sustained delivery of cargo, and mechanical tunability^29,47,48^. In addition, PNP hydrogels are easily administered due to their shear thinning and self-healing properties^46,49,50^. The fabrication process, which is easily scaled and therefore highly translatable, involves straightforward mixing of the polymer, nanoparticles (NPs), and an aqueous solution of cargo^47,51^. This combination of unique properties makes PNP hydrogels ideal candidates for use as a vaccine delivery platform.

Here, we report the use of PNP hydrogels as a vaccine delivery platform that affords simple encapsulation of vaccine components, provides sustained co-delivery of physicochemically distinct vaccine cargo, and creates a transient inflammatory niche upon administration for prolonged activation of the immune system. We show that these materials are easily injected and initiate the vaccine response locally through recruitment of antigen presenting cells (APCs), while also providing sustained release of vaccine cargo (Fig. 1). Importantly, our study shows an enhanced and prolonged humoral immune response to a single administration of vaccine-loaded PNP hydrogels, giving rise to average polyclonal antibody affinities equal to those of a monoclonal antibody. Overall, this study demonstrates the potential of the injectable PNP hydrogel platform to be used widely for improving subunit vaccine efficacy through simple encapsulation and prolonged delivery of these vaccines.

**Figure 1.**
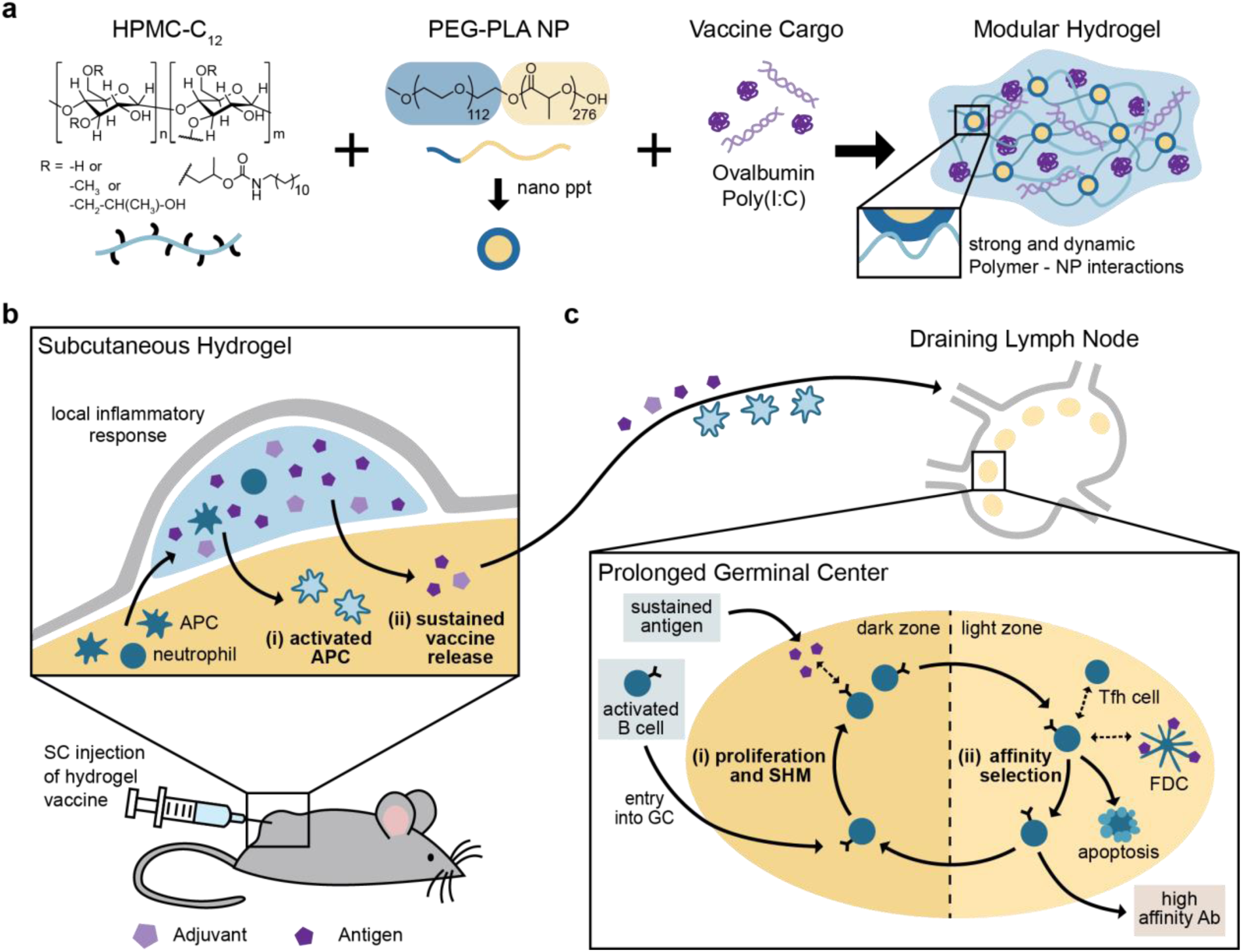
A schematic representation of the PNP hydrogel and proposed *in vivo* response to prolonged hydrogel-based vaccine delivery. **a**, Vaccine-loaded PNP hydrogels are formed when dodecyl-modified hydroxypropylmethylcellulose (HPMC-C_12_) is combined with poly(ethylene glycol)-*b*-poly(lactic acid) (PEG-PLA) nanoparticles and vaccine cargo, including ovalbumin (OVA) and Poly(I:C). Multivalent and dynamic non-covalent interactions between the polymer and nanoparticles constitute physical cross-links within the hydrogel structure. **b**, After subcutaneous (SC) injection of the hydrogel vaccine, local and migratory immune cell such as neutrophils and APCs infiltrate the gel, become activated, and then (**i**) activated APCs may migrate to the draining lymph nodes. The gel provides (**ii**) sustained release of the vaccine cargo to the draining lymph nodes, prolonging the germinal center response. **c**, The extended antigen availability in the germinal centers leads to increased (**i**) somatic hypermutation (SHM) and (**ii**) affinity selection, ultimately promoting higher affinity antibodies and a strong humoral immune response.

## Hydrogels for sustained vaccine exposure

We designed a PNP hydrogel material that can load vaccine components with high efficiency, is injectable, and can be tuned to co-deliver subunit vaccine components over prolonged timeframes. Our PNP hydrogels form rapidly when aqueous solutions of hydroxypropyl methylcellulose derivatives (HPMC-C_12_) are mixed with biodegradable polymeric NPs composed of poly(ethylene glycol)-*b*-poly(lactic acid) (PEG-PLA) (Fig. 1a, Table S1). Prior to mixing, the two components are solutions, but upon mixing a hydrogel rapidly forms multivalent and dynamic non-covalent interactions between the HPMC polymer and the PEG-PLA nanoparticles creating the physical cross-links that give rise to the hydrogel structure itself. Moreover, as is true with other supramolecular hydrogel platforms, all components that are mixed into the container become part of the self-assembled PNP hydrogel, ensuring 100% encapsulation efficiencies for any cargo. Furthermore, this simple synthesis enables the creation of numerous hydrogel formulations by changing the ratio of HPMC-C_12_ to NP to an aqueous solution^47^. For this manuscript, we chose two formulations based on their differences in mechanical properties: (i) 1 wt% HPMC-C_12_ + 5 wt% NP, designated “1:5”, and (ii) 2 wt% HPMC-C_12_ + 10 wt% NP, designated “2:10”. The future translation of our platform is simplified by the ease of synthesis along with the stability of the individual components of our system^52^.

The rheological properties of the hydrogel formulations of interest were then measured. Frequency-dependent oscillatory shear experiments, performed in the linear viscoelastic regime, demonstrated a formulation-dependent frequency response (Fig. 2a). At a representative angular frequency (ω = 10 rad/s), the 1:5 and 2:10 gels exhibited storage moduli (G’) of 50 Pa and 350 Pa, respectively. An angular frequency sweep showed that gels remained solid-like across a wide range of frequencies, with the G’ remaining above the loss modulus (G’’) at all frequencies tested (Fig. 2a, SI Fig. 1). A shear rate sweep showed that the viscosity of these PNP hydrogels decreased over two orders of magnitude with increasing shear rates, thereby demonstrating their ability to shear-thin (Fig. 2b, SI Fig. 1). The 1:5 gel exhibited a lower viscosity than the 2:10 gel across the shear rates tested: 10^−1^ to 10^2^ s^-1^ (Fig. 2b). The yield stress of these materials was determined by a stress ramp to be approximately 60 Pa and 300 Pa for the 1:5 and 2:10 gels, respectively (Fig. 2c, SI Fig. 2). These values describe both gels’ abilities to remain solid-like under low stresses (*i*.*e*., before and after injection), therefore maintaining a solid gel structure after injection and preventing flow from the injection site which facilitates the creation of a local inflammatory niche^50^.

**Figure 2.**
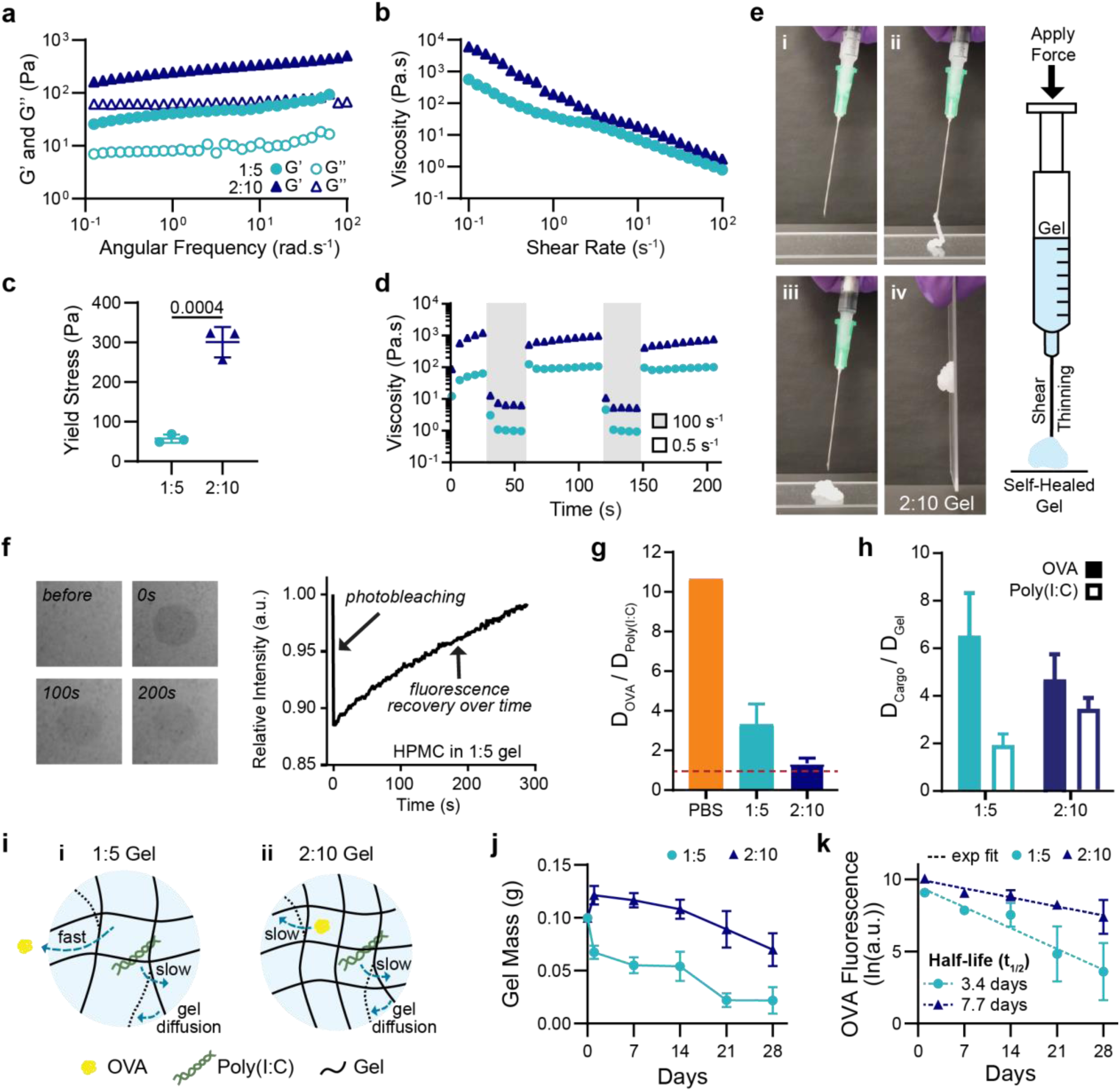
Material characterization and dynamics of entrapped molecular cargo. **a**, Frequency-dependent (σ = 1.8 Pa, 25°C) oscillatory shear rheology and **b**, steady shear rheology of two PNP hydrogel formulations designated “1:5” and “2:10”. **c**, Yield stress values from stress ramp measurements (n=3). **d**, Step-shear measurements of 1:5 and 2:10 gels over two cycles with alternating high shear (100 s-^1^) and low shear (0.05 s^-1^) rates. **e**, Images of 2:10 gel injection through a 21-gauge needle showing (**i**) before injection, (**ii**) during injection, (**iii-iv**) and after injection (screenshots from Video S1). **f**, FRAP experiment showing photobleaching of a select area at 0 sec, and the fluorescence recovering as fluorescent molecules diffuse back into the select area. **g**, Ratio of the diffusivity of OVA to the diffusivity of Poly(I:C) calculated from R_H_ values for PBS or using FRAP for the 1:5 and 2:10 gels. Values closer to one indicate more similar diffusivities of the OVA and Poly(I:C) (n=3). **h**, Ratio of the diffusivity of the cargo (OVA or Poly(I:C)) to the self-diffusivity of the hydrogel network. Values closer to one indicate that cargo diffusivity is limited by self-diffusion of the hydrogel (n=3). **i**, Representative schematic of (i) the 1:5 gel with OVA moving quickly and Poly(I:C) and the hydrogel matrix diffusing slower, and (ii) the 2:10 gel with the OVA, Poly(I:C), and hydrogel matrix all diffusing slowly. **j**, Gel mass over time following SC implantation measured from *in vivo* explants (n=5). **k**, Fluorescence of Alexa-647-OVA retained in explanted gels fit with an exponential decay to calculate t_1/2_ (n=5). All error bars are mean ± s.d., P value determined by two-tailed t-test.

Both gel formulations were shown to be injectable, a critical consideration for future translation of our material as a vaccine platform. Injectability was tested by measuring the viscosity of the gels when alternating between a high shear rate (100 s^-1^) reminiscent of the injection process, and a low shear rate (0.5 s^-1^) reminiscent of the working conditions upon implantation^45^. The viscosity of both gel formulations decreased by over two orders of magnitude with the applied high shear rate, and rapidly recovered (< 5 s) to their original viscosity when returned to a low shear rate (Fig. 2d and 2e, Video S1).

## Vaccine Cargo Dynamics

Subunit vaccines are composed of two main components: (i) antigen, which directs the antibody response to a specific substance, and (ii) adjuvant, which enhances the innate immune response^53^. These biomolecules can have vastly different sizes and physicochemical properties^53^, providing a challenge for controlled co-delivery because these characteristics typically dictate the diffusivity of these compounds and their release kinetics from hydrogel materials *in vitro* and *in vivo*^54^. To understand how PNP hydrogels can be used to deliver a wide range of cargo, we chose a model vaccine that incorporated an antigen and adjuvant with distinct physicochemical properties and very different molecular weights. We used the protein ovalbumin (OVA; MW = 43kDa) as a model antigen and Poly(I:C) (a toll-like receptor 3 agonist (TLR3); MW > 1MDa)^55,56^ as the adjuvant. Poly(I:C), a double-stranded RNA mimic that activates endosomal TLR3, is a potent type I interferon producer and has been used successfully as an adjuvant in preclinical vaccine studies, though it has not been approved for clinical use to date^56-58^. Importantly, this highly tunable materials platform enables these components to be easily replaced with other antigens and adjuvants for future applications.

To characterize the dynamics of vaccine exposure expected from the vaccine-loaded gels, we assessed the diffusivities of both vaccine cargo in each of the two gel formulations of interest using fluorescence recovery after photobleaching (FRAP) experiments (Fig. 2f, SI Fig. 3). The PNP gels are unique compared to traditional covalently cross-linked gels as their bonds are dynamic and continuously rearrange^47^. This feature results in self-diffusion of the polymer network over time within the intact hydrogel^59,60^. OVA’s hydrodynamic radius (R_H_) in PBS is reported in the literature to be 3.2 nm^61^, corresponding to a diffusivity value, D, of 76.6 μm^2^/s. Using multi-angle light scattering (MALS) we measured Poly(I:C)’s hydrodynamic radius to be much larger, with an R_H_ of 34.2 nm, corresponding to a D value of 7.2 μm^2^/s. This difference in hydrodynamic size results in OVA having more than a 10-fold greater diffusivity than Poly(I:C) in PBS (Fig. 2g) and a 100-fold difference in a typical covalently cross-linked hydrogel (SI Fig. 4). In contrast, FRAP measurements of the molecular cargo in the PNP gels demonstrated that the hydrogel mesh reduced cargo diffusivities by multiple orders of magnitude, whereby the diffusivity of OVA was determined to be 5.76 μm^2^/s in the 1:5 gel and 3.2 μm^2^/s in the 2:10 gel. The measured diffusivities for all experiments can be found in Table S2.

Interestingly, the ratio of OVA to Poly(I:C) diffusivities (D_OVA_/D_Poly(I:C)_) was 3.4 in the 1:5 gel, and only 1.4 in the 2:10 gel (Fig. 2g). When comparing the diffusivity of OVA to the self-diffusion of the hydrogel matrix, D_Gel_, which was determined by tracking the diffusivity of HPMC-C_12_ within the hydrogel, we saw that D_OVA_/D_Gel_ was ∼6.5 in the 1:5 gel, but only 4.7 in the 2:10 gel, while the ratio of Poly(I:C) to the hydrogel self-diffusion (D_Poly(I:C)_/D_Gel_) was approximately 2 for the 1:5 gel and 3.5 for the 2:10 gel (Fig. 2h). These data suggest that in both gels, Poly(I:C) is immobilized by the hydrogel mesh (*i*.*e*., Poly(I:C) is larger than the effective mesh size of the hydrogel; SI Fig. 4) such that the diffusion of Poly(I:C) in both gels is limited by the self-diffusion of the hydrogel network. The diffusion of OVA, however, is more limited by hydrogel self-diffusion in the 2:10 gel and appears to be small enough to diffuse more freely in the 1:5 gel (Fig. 2i, SI Fig. 4). Accordingly, the diffusivity of both OVA and Poly(I:C) is restricted by the self-diffusion of the hydrogel network in the 2:10 gel, enabling sustained co-delivery of the two cargo despite their large difference in physicochemical properties.

The *in vivo* cargo dynamics were characterized by assessing the timeframes of OVA retention in the subcutaneous (SC) space following *in vivo* administration. The 1:5 and 2:10 gels were loaded with Alexa Fluor 647 conjugated OVA and explanted to measure gel mass and cargo fluorescence at multiple time-points up to 4 weeks after vaccine administration. The 1:5 gels reached half of their initial mass after 2 weeks whereas the 2:10 gels retained half of their initial mass past 4 weeks (Fig. 2j). The cargo fluorescence was measured after explantation and mechanical disruption. From these *in vivo* data we calculated an OVA retention half-life of 3.4 days and 7.7 days for the 1:5 and 2:10 gels, respectively (Fig. 2k).

## Humoral immune response to vaccination

To investigate if sustained vaccine exposure from a single administration of the gel-based vaccines (*i*.*e*., gel carrying OVA and Poly(I:C)) could enhance the magnitude and duration of the humoral response, we quantified antigen specific antibodies in the serum after SC injection (Fig. 3a). Humoral immunity is a component of the adaptive immune response mediated by antibodies, which are critical to viral immunity^53^. The peak concentration of OVA-specific serum antibodies was 2-3 times higher for mice receiving the gel-based vaccines than mice receiving the same vaccine in a standard PBS bolus administration (Fig. 3b). The 1:5 and 2:10 gel-based vaccines yielded antibody concentrations higher than the bolus peak response past day 90 (Fig. 3b). We also observed that the gel alone acts as a mild adjuvant, but the addition of Poly(I:C) significantly improves the potency and durability of the antibody response (SI Fig. 5). Additionally, delivering antigen and adjuvant in separate gels on contralateral flanks was not as effective as co-delivery of OVA and Poly(I:C) from the same gel (SI Fig. 5). The increased magnitude and duration of the primary antibody response suggests that delivery of vaccines in PNP hydrogels would afford more durable protection against target pathogens than bolus administration of the same vaccines.

**Figure 3.**
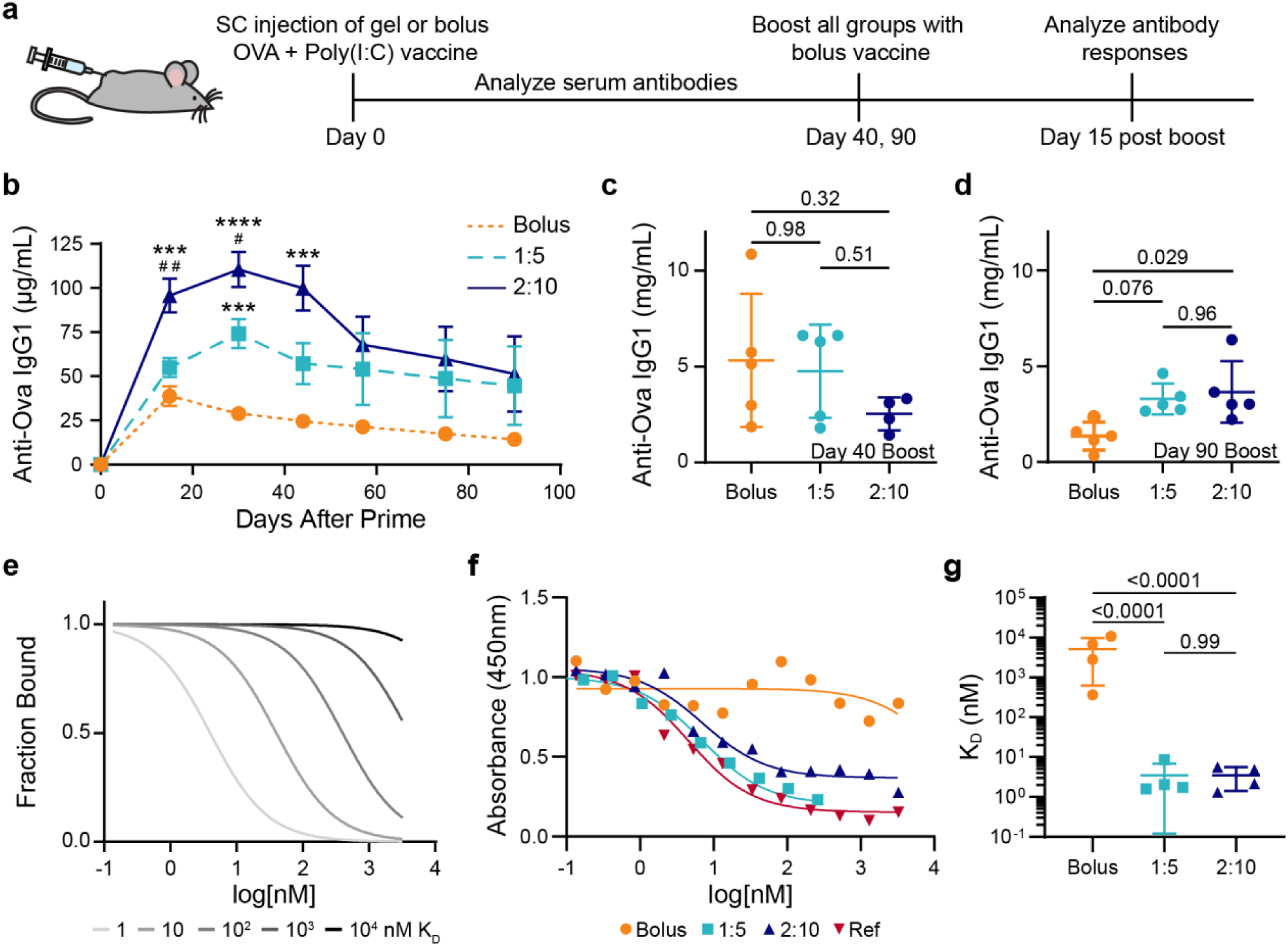
Antibody concentration and affinity following immunization. **a**, Timeline of the experimental setup shows SC injection of a model vaccine containing OVA and Poly(I:C) in a gel or bolus formulation at day 0, antibody analysis over time following a single administration, boost with a bolus vaccine formulation at day 40 or 90, and analysis of the immune response 15 days after the boost. **b**, Serum anti-OVA IgG1 concentrations from day 0 to day 90 after a single injection of vaccines (n = 5 to 19; 1 to 4 independent experiments; mean ± s.e.m.). ***p < 0.001 and ****p < 0.0001 compared to bolus, ^#^p < 0.05, ^# #^p < 0.005 compared to 1:5, determined by mixed-effects analysis with Tukey’s post hoc test. **c** and **d**, Serum anti-OVA IgG1 concentrations 15 days after bolus boost on day (**c**) 40 or (**d**) 90 for animals receiving either bolus, 1:5 gel, and 2:10 gel vaccines (n = 4 to 5; mean ± s.d.). Reported P values determined by one-way ANOVA with Tukey’s post hoc test. **e**, Model comparing competitive binding data with K_D_ ranging from 1 to 10^4^ nM. **f**, Representative competitive binding curves for bolus, 1:5 gel, and 2:10 gel vaccine groups after the day 90 boost compared to a mAb reference competing with the same mAb. **g**, Calculated K_D_ values from fitted binding curves for 2:10 gel and bolus vaccine groups (n = 4; mean ± s.d.). P values determined by one-way ANOVA with Tukey’s post hoc test.

Most vaccines are administered with booster shots to increase the immune memory and ensure long-term protective antibodies are produced^53^. To study the prime-boost response, we administered OVA and Poly(I:C) in PBS for all groups at 40 or 90 days post-prime in separate mouse cohorts (Fig. 3a). The OVA-specific serum IgG1 antibodies were assessed 15 days after the boost^62^. The day 40 boost led to statistically equivalent antibody concentrations in all groups (Fig. 3c); however, when the boost was administered 90 days after priming, the mice primed with the 2:10 gel responded with higher antibody titers than the bolus vaccine primed mice (Fig. 3d). Both gel groups exhibited an equivalently strong antibody response to boosts administered at both time-points (Fig. 3c and 3d), indicating robust humoral memory compared to the bolus group, which had a 3.8-fold decrease in antibody production in response to boosts at 40 days and 90 days (p=0.038). The described antibody responses all refer to IgG1 antibodies, which were the most highly represented among the IgG subclasses. Characterization of IgG2b and IgG2c anti-OVA antibodies can be found in the supplemental information (SI Fig. 6). Importantly, we see no detectable anti-PEG antibodies developed and no noticeable fibrotic response to these hydrogel-based vaccines (SI Fig. 7, SI Fig. 8).

To investigate the quality of the humoral response, we also determined the affinities of the serum antibodies after the 90-day boost towards OVA using a competitive binding enzyme-linked immunosorbent assay (ELISA) (Fig. 3e-g). We found that the polyclonal population of antibodies produced by the mice that received the 1:5 or 2:10 gel vaccine prime had K_D_ values of approximately 3 nM, while the antibodies produced by the mice receiving OVA and Poly(I:C) in a PBS bolus prime had a K_D_ value of 3,000 nM (Fig. 3f and 3g, SI Fig. 9). PNP gel-based vaccination, therefore, results in at least a 1000-fold enhancement in affinity maturation of the antibodies. We validated the trends seen in the competitive binding ELISA with surface plasmon resonance (SPR) experiments (SI Fig. 10). The increase in antibody affinity observed for the gel groups suggests that the prolonged vaccine exposure with the gel prime led to an enhancement of GC responses and B cell affinity selection^8^.

## Local inflammatory niche within the hydrogel depot

We evaluated the cells infiltrating the gel depots to understand if the gel was creating an inflammatory niche in addition to providing sustained release of the cargo (Fig. 4a and 4b). We quantified cell infiltration into the OVA and Poly(I:C) vaccine-loaded 2:10 gel compared to the 2:10 gel alone 7 days after SC injection *in vivo* with flow cytometry (SI Fig. 11). We found that the presence of vaccine cargo increased total cell infiltration into the gels, whereby almost 1.0×10^6^ total cells were found within a vaccine-loaded gel compared to 0.2×10^6^ cells in the control gel (Fig. 4c). The vaccine-loaded gel did not significantly recruit more neutrophils, but did recruit significantly higher numbers of monocytes, macrophages, and dendritic cells than an control gel (Fig. 4d-g)^63^. Within the dendritic cell population found in the vaccine-loaded gel, the majority were migratory type 2 conventional DCs (cDC2), which play a critical role in activating follicular T helper cells (Tfh) and initiating the humoral immune response (Fig. 4h)^64^. OVA uptake by cDC2s was confirmed via flow cytometry using Alexa Fluor 647 conjugated OVA as part of the vaccine (Fig. 4i). Diverse cell populations were observed in both the vaccine loaded gel and control gel, including neutrophils, monocytes, macrophages, dendritic cells, as well as other myeloid and non-myeloid cells (Fig. 4j). The initiation of the adaptive immune response relies on signaling from APCs, therefore the abundance of APCs (*i*.*e*., macrophages and DCs) in the vaccine-loaded gel at this early timepoint suggests that the gel acts as a beneficial inflammatory niche for immune activation.

**Figure 4.**
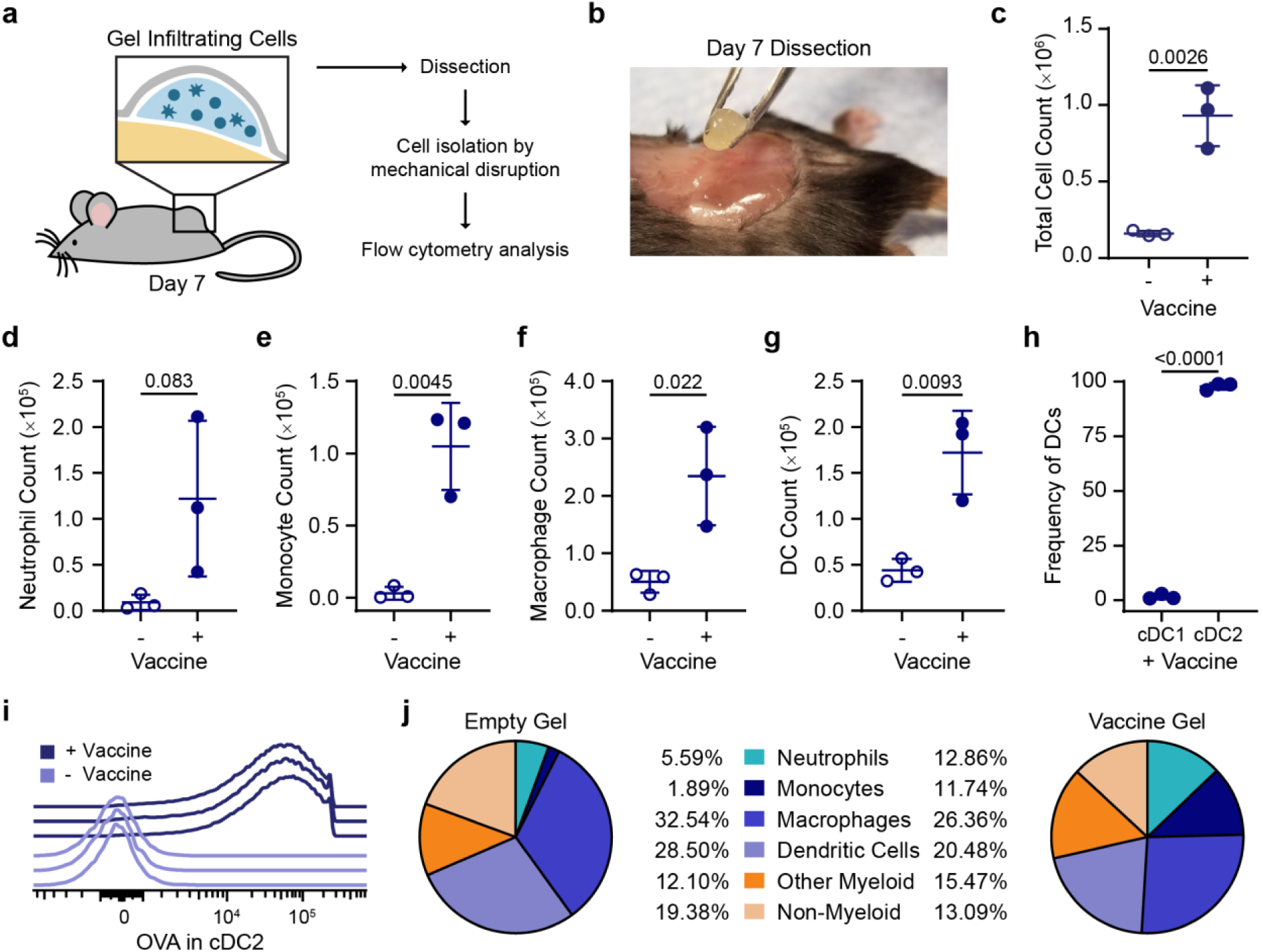
Characterization of the local inflammatory niche. **a**, Schematic of the inflammatory niche within the gel depot and an experimental flow chart. **b**, Picture of surgical removal of 2:10 gel after 7 days *in vivo*. **c**, Total cells in 2:10 gel with or without vaccine (OVA+Poly(I:C)) were quantified using flow cytometry. **d to g**, Total count of neutrophils (**d**), monocytes (**e**), macrophages (**f**), and DCs (**g**) found in the empty and vaccine-loaded 2:10 gels. **h**, The frequency of cDC1 (XCR1^hi^CD11b^lo^) and cDC2 (XCR1^lo^CD11b^hi^) of the total dendritic cells (DCs) in the vaccine-loaded 2:10 gel. **i**, Histogram of the Alexa-647 OVA signal in cDC2s from individual mice with and without the vaccine. **j**, The frequency of neutrophils, monocytes, macrophages, DCs, other myeloid cells, and non-myeloid cells within the CD45^+^ cell populations found in the empty and vaccine-loaded 2:10 gels. For **c-j**, n=3 mice. All error bars are mean ± s.d., P values determined by two-tailed t-test.

## Germinal center response to vaccination

We hypothesized that the enhanced humoral response observed for gel-based sustained-exposure vaccines was due to the prolonged lifetime of GCs in the lymph nodes. GCs are dynamic sites that form after the activation of germinal center B cells (GCBCs) and are responsible for producing memory B cells and high-affinity antibodies^53^. The GCs of the 2:10 and bolus groups 15 days after vaccination with OVA and Poly(I:C) were qualitatively visualized with immunohistochemistry (Fig. 5a). To quantitatively evaluate the GC response, we measured the frequency of GCBCs in the draining lymph nodes 15 and 30 days after vaccination with OVA and Poly(I:C) (SI Fig. 12). At 15 days after vaccine administration, mice in both the 1:5 and 2:10 gel groups had significantly higher frequencies of GCBCs than the mice who received the vaccine in PBS (Fig. 5b). At 30 days post-vaccination, mice that received the 2:10 gel formulation continued to have a higher GCBC frequency (Fig. 5c). Moreover, the 1:5 and 2:10 gel groups had a higher frequency of class-switched GCBCs (IgG1^+^) compared to the bolus group at day 15 (Fig. 5d); this is a critical indicator of a protective humoral respnonse^53,65^. At day 30, the percent of class-switched GCBCs remained higher for the 2:10 group compared to the vaccine in PBS (Fig. 5e). Tfh cells play a critical role in GCBC selection, class-switching, and differentiation, and GCBCs interact with Tfh cells mainly in the light zone (LZ) of the GC (Fig. 5f)^66^. In agreement with the GCBC results, we found an increase in the frequency of Tfh cells in the lymph nodes of both gel groups compared to bolus administration at the 15-day timepoint (Fig. 5g). The gel also led to an increase in the ratio of light zone to dark zone (LZ/DZ) GCBCs compared to bolus administration (Fig. 5h). These data indicate an increase in affinity selection compared to expansion and somatic hypermutation^53^. Together, these data suggest that the prolonged vaccine exposure from the gels enhanced the magnitude and duration of the GC response and antibody affinity maturation.

**Figure 5.**
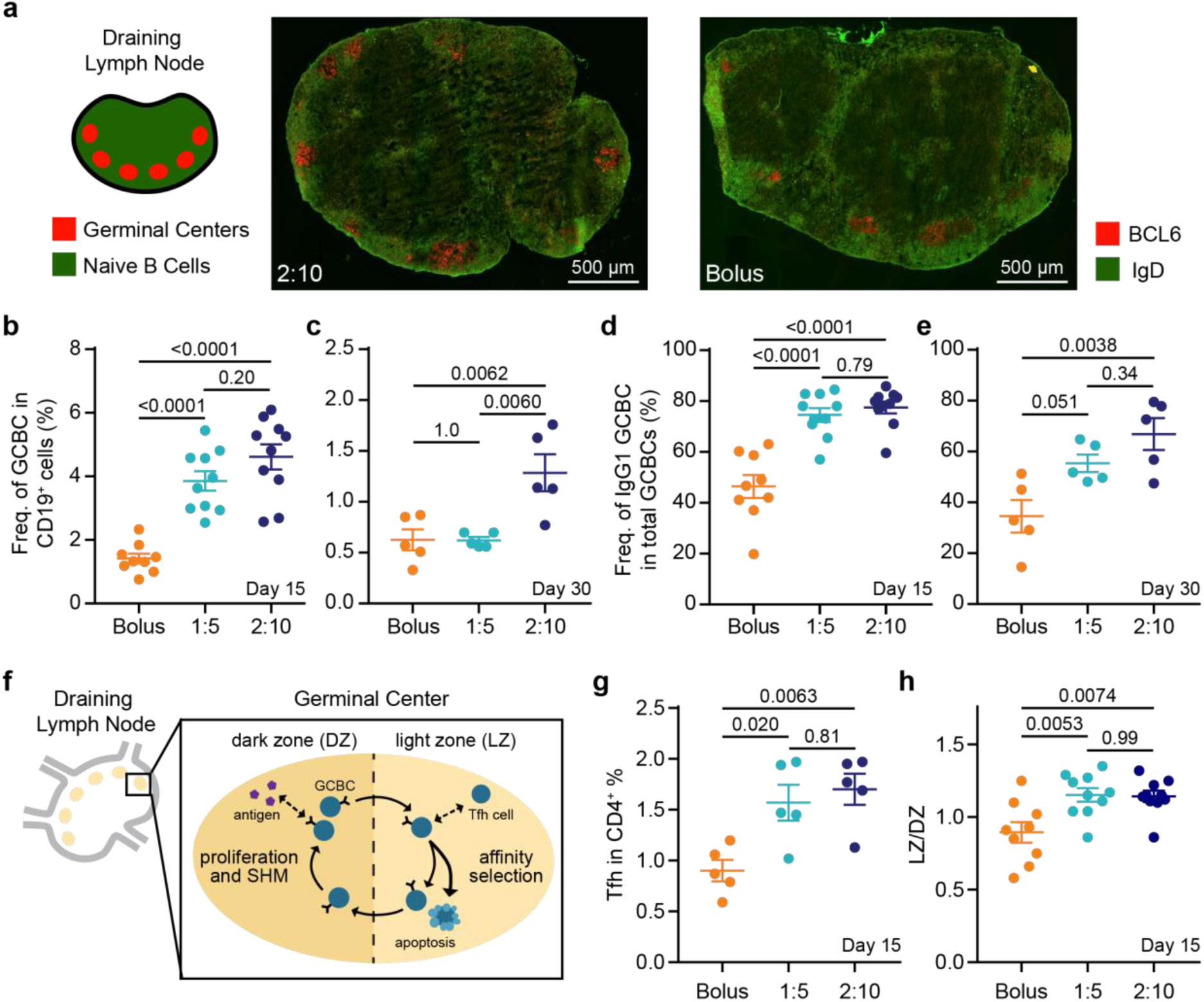
Germinal center response to single vaccine administration. **a**, Immunohistochemistry (IHC) of explanted inguinal lymph node 15 days after OVA+Poly(I:C) vaccine administration in 2:10 and bolus groups to visualize germinal centers (red) and naïve B cells (green). **b and c**, The frequency of GCBCs within total B cells at day 15 (**b**) and day 30 (**c**) after prime and **d and e**, frequency of IgG1+ GCBCs within total GCBCs at day 15 (**d**) and day 30 (**e**) after prime in the inguinal lymph nodes were measured by flow cytometry (n = 5 to 10). **f**, Schematic of the GC response. **g**, The percent of Tfh cells out of the CD4^+^ cell population and **h**, and the ratio of LZ to DZ GCBCs in the inguinal lymph nodes at day 15 after vaccinatation (n = 5 to 10). For **b,d**, and **h** data come from 2 independent experiments, all other graphs represent 1 independent experiment. All error bars are mean ± s.d., P values determined by one-way ANOVA with Tukey’s post hoc test.

## Discussion

In this study, we designed an injectable hydrogel material capable of prolonged co-delivery of physicochemically-distinct cargo on timescales relevant to the kinetics of antigen exposure typical of natural infections. The vaccine-loaded PNP hydrogels are injectable and retain their solid-like structure when under low stresses, enabling creation of a new stimulatory microenvironment within the body while facilitating sustained vaccine delivery. Additionally, the dynamic networks comprising the 1:5 and 2:10 gels provide unique cargo diffusion characteristics compared to traditional covalent hydrogel systems. Our observations suggest that the 2:10 gel mesh sufficiently constrains the diffusion of both OVA and Poly(I:C), even though Poly(I:C) is much larger than OVA, such that their diffusion and release is dictated by the self-diffusion of the dynamically cross-linked hydrogel network. These physical properties ensure co-presentation of antigen and adjuvant over prolonged timeframes to the immune system which has been a long-standing challenge within the field.

Developing effective humoral immune responses towards difficult pathogens, such as HIV-1 or malaria, requires the creation of high affinity antibodies^67,68^. The sustained vaccine exposure enabled by these gels was shown to increase the magnitude and duration of GC responses in the draining lymph nodes, leading to enhancements in the magnitude, persistence, and quality of the humoral immune response against the antigen. The sustained release of antigen from PNP hydrogels is able to better mimic natural infections where GCBCs continuously receive antigen-derived signaling to undergo many rounds of selection and somatic hypermutation (Fig. 1c). In addition, sustained vaccine release provides GCBCs the essential B cell receptor signaling needed to promote the DZ to LZ transition and positive selection of high affinity GCBCs^6^. The significantly higher affinity antibodies produced in mice immunized with gel-based vaccines suggest an increase in somatic hypermutation and commensurate enhancement in affinity maturation.

These gels also act as a local stimulatory microenvironment where infiltrating cells experience high local concentrations of adjuvant and antigen. Implantable biomaterials have been shown to act as immune-stimulating niches which recruit and activate APCs that then migrate to the lymph nodes^69^. Herein, we observed minimal immune cell infiltration in the gels alone; however, loading the gels with vaccine cargo dramatically increased immune cell infiltration, which is likely driven by the inflammatory signals initiated by the entrapped adjuvant. The vaccine-loaded gel was able to recruit macrophages and DCs, which are professional APCs that are essential for initiating the adaptive immune response. In particular, almost all infiltrating DCs were found to be cDC2s which are known to play a critical role in activating Tfh cells and initiating the humoral immune response. We hypothesize that these cells are activated by entrapped adjuvant and migrate to the draining lymph nodes. The draining lymph nodes are therefore receiving activated immune cells from the gel as well as the soluble vaccine cargo as it is released from the gel depot over time (Fig. 1b and 1c).

In conclusion, PNP hydrogels provide a simple and effective platform for sustained delivery of subunit vaccines to increase the potency and durability of the humoral immune response. Using our platform as a tool to probe the interactions between the immune system and a vaccine depot will enable more precise material development for vaccine delivery. Our PNP hydrogel represents a highly tunable platform for effectively manipulating the humoral immune response for any subunit vaccine of interest.

## Materials and Methods

### Materials

HPMC (meets USP testing specifications), N,N-Diisopropylethylamine (Hunig’s base), hexanes, diethyl ether, N-methyl-2-pyrrolidone (NMP), dichloromethane (DCM), lactide (LA), 1-dodecylisocynate, and diazobicylcoundecene (DBU) were purchased from Sigma Aldrich and used as received. Monomethoxy-PEG (5 kDa) was purchased from Sigma Aldrich and was purified by azeotropic distillation with toluene prior to use.

### Preparation of HPMC-C_12_

HPMC-C_12_ was prepared according to previously reported procedures^47^. HPMC (1.0 g) was dissolved in NMP (40 mL) by stirring at 80 °C for 1 h. Once the solution cooled to room temperature, 1-dodecylisocynate (105 mg, 0.5 mmol) and N,N-Diisopropylethylamine (catalyst, ∼3 drops) were dissolved in NMP (5.0 mL). This solution was added dropwise to the reaction mixture, which was then stirred at room temperature for 16 h. For rhodamine conjugated HPMC-C_12_, 10 mg of rhodamine B isothiocyanate (0.019 mmol) was added to the 1-dodecylisocyanate solution before adding it dropwise to the reaction mixture. This solution was then precipitated from acetone, decanted, re-dissolved in water (∼2 wt%), and placed in a dialysis tube for dialysis for 3-4 days. The polymer was lyophilized and reconstituted to a 60 mg/mL solution with sterile PBS.

### Preparation of PEG-PLA NPs

PEG-PLA was prepared as previously reported^47^. Monomethoxy-PEG (5 kDa; 0.25 g, 4.1 mmol) and DBU (15μL, 0.1 mmol; 1.4 mol% relative to LA) were dissolved in anhydrous dichloromethane (1.0 mL). LA (1.0 g, 6.9 mmol) was dissolved in anhydrous DCM (3.0 mL) with mild heating. The LA solution was added rapidly to the PEG/DBU solution and was allowed to stir for 10 min. The reaction mixture was quenched and precipitated by 1:1 hexane and ethyl ether solution. The synthesized PEG-PLA was collected and dried under vacuum. Gel permeation chromatography (GPC) was used to verify that the molecular weight and dispersity of polymers meet our quality control (QC) parameters.

NPs were prepared as previously reported^47^. A 1 mL solution of PEG-PLA in DMSO (50 mg/ml) was added dropwise to 10 mL of water at room temperature under a high stir rate (600 rpm). NPs were purified by ultracentrifugation over a filter (molecular weight cut-off of 10kDa; Millipore Amicon Ultra-15) followed by resuspension in water to a final concentration of 200 mg/mL. NPs were characterized by dynamic light scattering (DLS) to find the NP diameter, 32 ± 4 nm, and zeta potential, -28 ± 7 mV (SI Table 2).

### PNP Hydrogel Preparation

The 2:10 formulation contained 2 wt% HPMC-C_12_ and 10 wt% PEG-PLA NPs in PBS. These gels were made by mixing a 2:3:1 weight ratio of 6 wt% HPMC-C_12_ polymer solution, 20 wt% NP solution, and PBS. The 1:5 formulation contained 1 wt% HPMC-C_12_ and 5 wt% PEG-PLA NPs in PBS. These gels were made by mixing a 2:3:7 weight ratio of 6 wt% HPMC-C_12_ polymer solution, 20 wt% NP solution, and PBS. The solutions were mixed with a spatula, mildly centrifuged to remove bubbles arising from mixing, and then loaded into a syringe.

### Vaccine Formulations

The model vaccine contained a 100 μg dose of OVA (Sigma Aldrich) and 50 μg dose of Poly(I:C) (Sigma Aldrich) per 100 μL of gel or PBS. For the bolus vaccines, the above vaccine concentrations were prepared in PBS and loaded into a syringe for administration. For the PNP hydrogels, the vaccine cargo was added at the appropriate concentration into the PBS component of the gel before adding the polymer and NP solutions, as described above.

### Material Characterization

Rheological characterization was performed using a TA Instruments Discovery HR-2 torque-controlled rheometer fitted with a Peltier stage. All measurements were performed using a serrated 20-mm plate geometry at 25 °C.

Dynamic oscillatory frequency sweep measurements were performed with a constant torque (2 uN.m; σ = 1.27 Pa) from 0.1 rad/s to 100 rad/s. Steady shear experiments were performed from 0.1 to 100 s^-1^. Step-shear experiments were performed by alternating between a low shear rate (0.05 s^-1^) and high shear rate (100 s^-1^) for two full cycles.

### FRAP Analysis

Alexa Fluor 647 conjugated OVA (Thermo Fisher Scientific), rhodamine conjugated Poly(I:C) (Invivogen), and rhodamine conjugated HPMC-C_12_ were used to visualize the diffusion of the cargo and gel. Samples were photobleached with a 50 µm diameter for the region of interest (ROI). Different tests (n = 5) were made for 3 different samples from the same batch at different locations of the sample. A spot was bleached with a pixel dwell time of 177.32 μs. 500 post-bleach frames were recorded at 1 frame/s to form the recovery exponential curve. The diffusion coefficient was calculated as^70^:

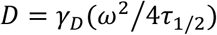

Where the constant *γ*_*D* =_ *τ*_1/2_/*τ*_*D*_, with *τ*_1/2_ being the half-time of the recovery, *τ*_*D*_ the characteristic diffusion time, both yielded by the ZEN software, and *ω* the radius of the bleached ROI (25 µm).

The diffusivity of cargo in PBS was calculated using the Stokes-Einstein Law Equation for diffusion^71^ where k_B_ is Boltzmann’s constant, T is temperature in Kelvin, η is solvent viscosity, and R is solute hydrodynamic radius:

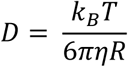

The diffusivity of cargo in a model covalent PEG gel was calculated using the Multiscale Diffusion Model (MSDM) assuming 25 °C, 5% volume fraction, and 35 nm mesh size^72^. The calculated values are comparable to experiment diffusivities of similar sized cargo found in the literature^73^.

### Mice and Vaccination

C57BL/6 (B6) mice were purchased from Charles River and housed at Stanford University. Female mice between 6 and 10 weeks of age at the start of the experiment were used. The mice were shaved several days before vaccine administration and received a subcutaneous injection of 100 μL gel or bolus vaccine on their backs under brief isoflurane anesthesia. PBS injections used a 26-gauge needle, and gel injections used a 21-gauge needle. Mouse blood was collected from the cheek or tail vein for survival studies, or through cardiac puncture for terminal studies. The inguinal LN’s and SC gels were collected for GC and cell infiltration analysis after euthanasia.

### In Vivo Cargo Dynamics

Vaccine loaded gels with Alexa Fluor 647 conjugated OVA (Thermo Fisher Scientific), were explanted at 1, 7, 14, 21, or 28 days, and weighed to report the gel erosion over time. After weighing, the explanted gels were diluted in PBS and homogenized using a glass dounce homogenizer (Wheaton). The fluorescence of the homogenized gels was read with ex: 650 nm and em: 665 nm on a plate reader (Tecan Infinite M1000). Raw fluorescence values were normalized for polymer background, the natural log of these values was plotted and fit with linear equations to find the rate constant using GraphPad Prism 7.04 (GraphPad Software). This rate constant was used to calculate a half-life of cargo retention.

### Antibody Concentration and Affinity

Serum antibody concentrations and affinity for the OVA model vaccine were measured using an anti-ovalbumin mouse IgG1 ELISA (Cayman Chemicals, 500830). For time course measurements the serum was diluted 1:1,000 in assay buffer and for post-challenge analysis the serum was diluted 1:100,000. The assay was performed according to the manufacturer’s instructions to find concentration. The plates were analyzed using a Synergy™ H1 Microplate Reader (BioTek Instruments) at 450nm. Serum antibody concentrations were calculated from the standard curves and represented as μg/mL or mg/mL.

Anti-OVA IgG2b and IgG2c antibody concentrations were measured using mouse anti-OVA antibody assay kits for IgG2b (chondrex, 3016) and IgG2c (Chondrex,3029). Serum was diluted 1:1,000 and the assay was performed according to the manufacturer’s instructions to find concentration. The plates were analyzed using a Synergy™ H1 Microplate Reader (BioTek Instruments) at 450nm. Serum antibody concentrations were calculated from the standard curves and represented as μg/mL.

For affinity, dilutions of the serum were mixed with a constant 3 nM of an HRP conjugated anti-OVA antibody (BioLegend) and incubated for 2 h at room temperature. The wells were washed, incubated with TMB substrate (TMB ELISA Substrate (High Sensitivity), Abcam), and the reaction was stopped with 1 M HCl. Absorbance was read with a plate reader at 450 nm. Data was fit with a one-site competitive binding model with Graphpad Prism. The control antibody was assumed to have a K_D_ of 1 nM based on common affinities of mAbs found in the industry. This assumption affects only the absolute K_D_ values reported and not the relative differences between treatment groups. Statistics were performed on the log_10_(K_D_) values. Individual binding curves can be found in Supplementary Fig. 3.

### Immunohistochemistry in LN

Draining inguinal lymph nodes were isolated and frozen in molds containing OCT medium on dry ice. Frozen lymph nodes were sectioned at 7 µm and stored at −80 °C. Sections were stained with biotin labeled anti mouse IgD (eBioscience) and PE labeled anti mouse BCL6 (BD Biosciences) antibodies, and subsequently stained with Al488 conjugated Streptavidin (Thermo Fisher Scientific). Fluorescent images were captured using a 20X objectives on a fluorescence microscope (Keyence). There were fundamental limitations in the quantity and orientation of GCs in the LNs, so immunohistochemistry was not used to quantify cell types or GC features.

### Immunophenotyping in LN

Inguinal lymph nodes were surgically removed from the animals after euthanasia and then were mechanically disrupted to create a cell suspension. For FACS analysis, cells were blocked with Fc receptor antibody (clone: 2.4G2, Tonbo Biosciences) and then stained with fluorochrome conjugated antibodies: CD19, GL7, CD95, CXCR4, CD86, IgG1, CD4, CXCR5, and PD1. After staining, cells were washed and fixed with 1.5% paraformaldehyde (PFA). Stained cells were analyzed on LSRII flow cytometer. Data were analyzed with FlowJo 10 (FlowJo LLC). See SI Fig. 11 for gating strategy and SI Table 3 for the antibody panel.

### Immunophenotyping in Gel

Gels were surgically removed from the animals after euthanasia. The gels were processed using mechanical disruption to create a cell suspension which was then filtered through a 70 μm cell strainer (Celltreat). To count total cells in the gels, the single-cell suspensions were stained with anti-mouse CD45 and DAPI to count live leukocytes. The cells were mixed with CountBright Absolute Counting beads (ThermoFisher) and analyzed on a LSRFortessa X-20 flow cytometer (BD Biosciences). To analyze neutrophil and other myeloid cell counts in the gels, the single-cell suspensions were incubated with anti-CD16/CD32 (produced in house from 2.4G2 hybridoma) to block Fc receptor binding, and then stained with anti-mouse CD45, CD3, CD19, CD11c, Ly6G, CD11b, and MHCII in FACS buffer (2mM EDTA, 2% FBS in PBS) for 30 min on ice. Dead cells were excluded by DAPI staining. Cells were acquired on a LSRFortessa X-20 and fcs files were analyzed using FlowJo 10 (FlowJo LLC). See SI Fig. 10 for gating strategy and SI Table 3 for the antibody panel.

### Animal Protocol

All animal studies were performed in accordance with National Institutes of Health guidelines, with the approval of Stanford Administrative Panel on Laboratory Animal Care.

### Statistical Analysis

All results are expressed as a mean ± standard deviation (s.d.) except for figure 3b which is mean ± standard error of the mean (s.e.m.) as indicated in the figure captions. Comparisons between two groups were conducted by a two-tailed Student’s t-test. One-way ANOVA test with a Tukey’s multiple comparisons test was used for comparison across multiple groups. Statistical analysis was run using GraphPad Prism 7.04 (GraphPad Software). Statistical significance was considered as p < 0.05.

## Data Availability

The data that support the findings of this study are available from the corresponding author upon reasonable request.

## Acknowledgements

This research was financially supported by the Center for Human Systems Immunology with Bill and Melinda Gates Foundation (OPP1113682) and the ITI seed grant. G.A.R. is grateful for the NSF Graduate Research Fellowship (DGE-114747). A part of this work was performed at the Stanford Nano Shared Facilities (SNSF), supported by the National Science Foundation under award ECCS-1542152, and the Stanford High Throughput Bioscience Center (HTBC).

