## Supplemental Information for "Injectable hydrogels for sustained co-delivery of subunit vaccines enhance humoral immunity"

### Supplementary Information

#### Supplemental Figures

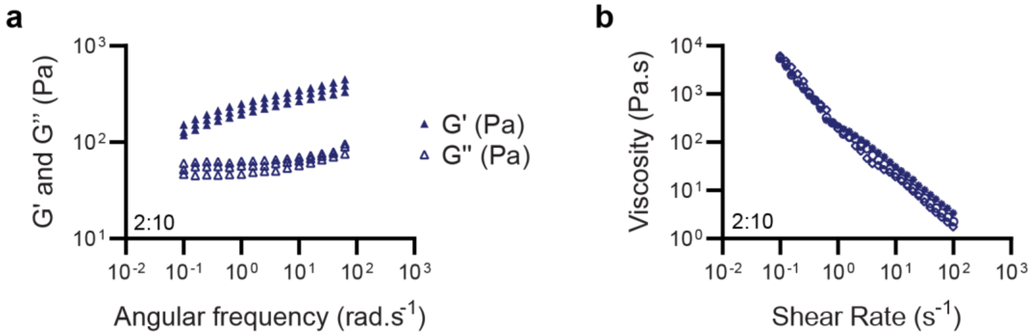

**Figure S1 | Batch to batch consistency of rheological characterization. a,** Replicates of frequency-dependent (σ = 1.8 Pa, 25 °C) oscillatory shear rheology and **b**, steady shear rheology of 2:10 gels (n=3) demonstrating batch to batch consistency.

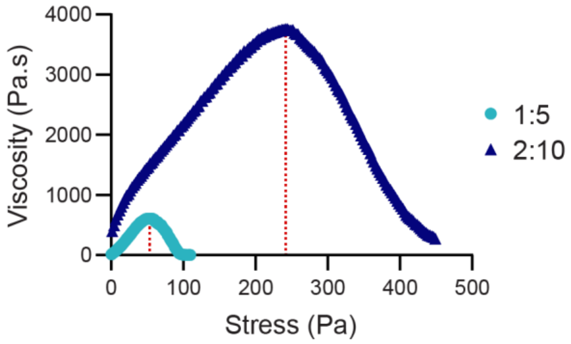

**Figure S2 | Raw data for yield stress determination**. Representative stress ramp rheological experiments for 1:5 and 2:10 gels with the peak viscosity indicated by a red line showing how the yield stress was measured.

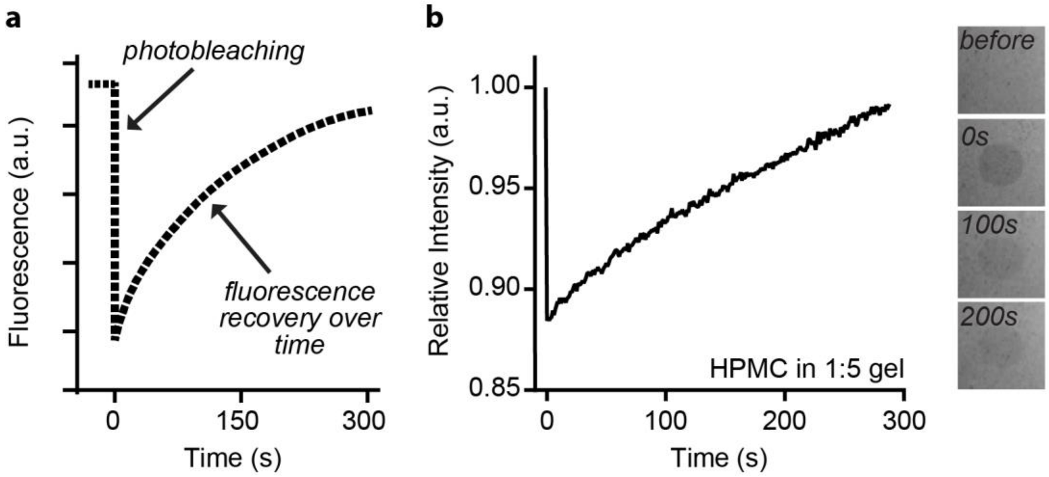

**Figure S3 | A representative fluorescence recovery after photobleaching (FRAP) experiment**. FRAP was used to characterize the mobility of the cargo and polymer through the hydrogel systems**. a,** Several frames using a low light level are acquired to determine the initial fluorescence, and then a high intensity of light is applied for a short time inside a region of interest to bleach the fluorescence in the sample. Finally, the recovery of fluorescence is monitored to measure how fast the molecule of interest redistributes. **b,** The figure shows raw data probing the self-diffusion of HPMC within the weak hydrogels.

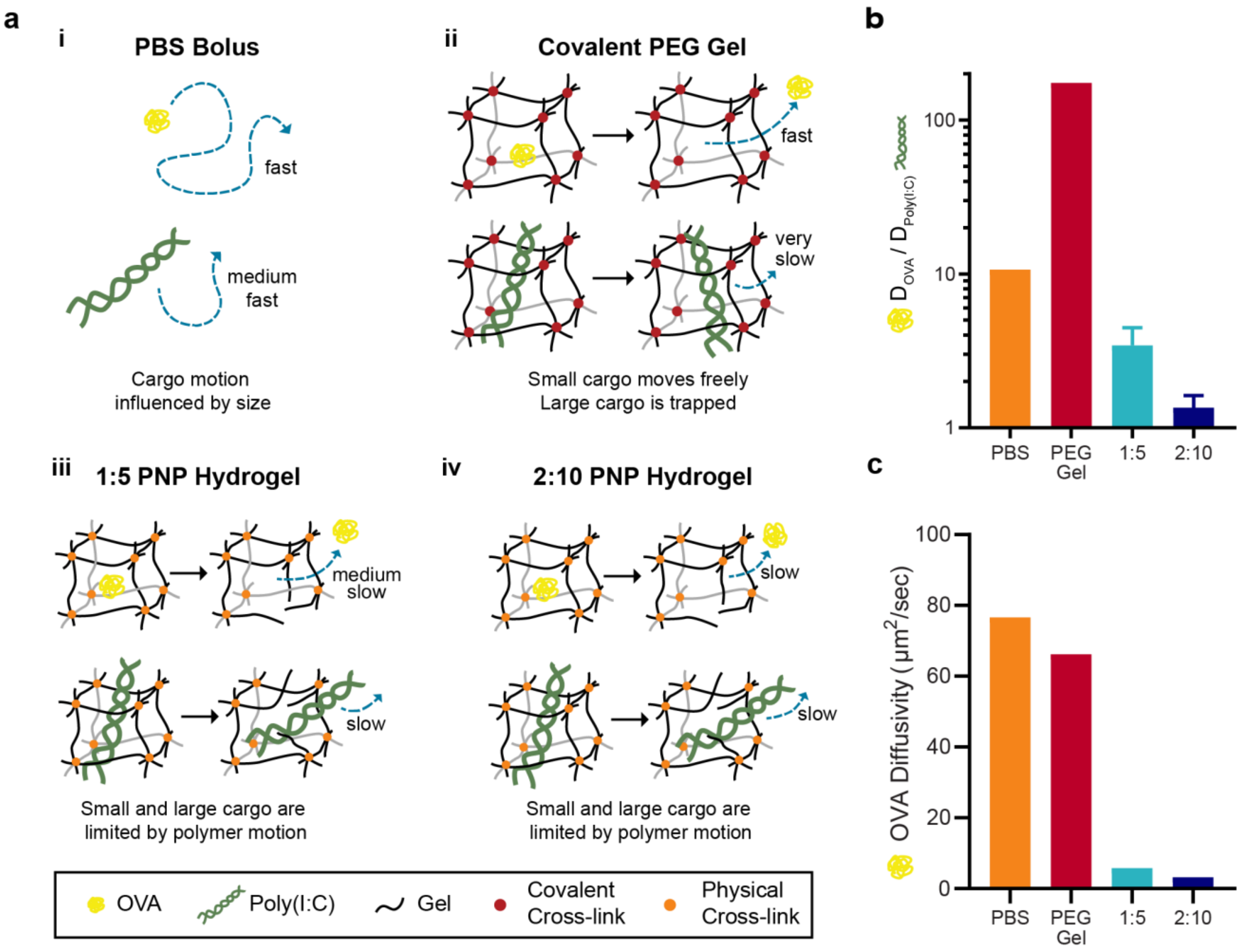

**Figure S4 | Cargo diffusivity schematics and values**. **a**, Representations of the diffusivity of OVA (43kDa) and Poly(I:C) (>1MDa) in (i) PBS, (ii) a covalently crosslinked PEG hydrogel, (iii) 1:5 PNP gel, and (iv) 2:10 PNP gel. **b**, The ratio of the diffusivity of OVA to the diffusivity of Poly(I:C), where values closer to one indicate more similar diffusivities, and **c**, absolute OVA diffusivities in each matrix. PBS diffusivities were calculated from $R_H$ values using the Stokes-Einstein equation, and PEG diffusivities were calculated with $R_H$ values using a multiscale diffusion model[1], while the 1:5 and 2:10 gel diffusivities were determined using FRAP experiments described in Figure 2 of the manuscript.

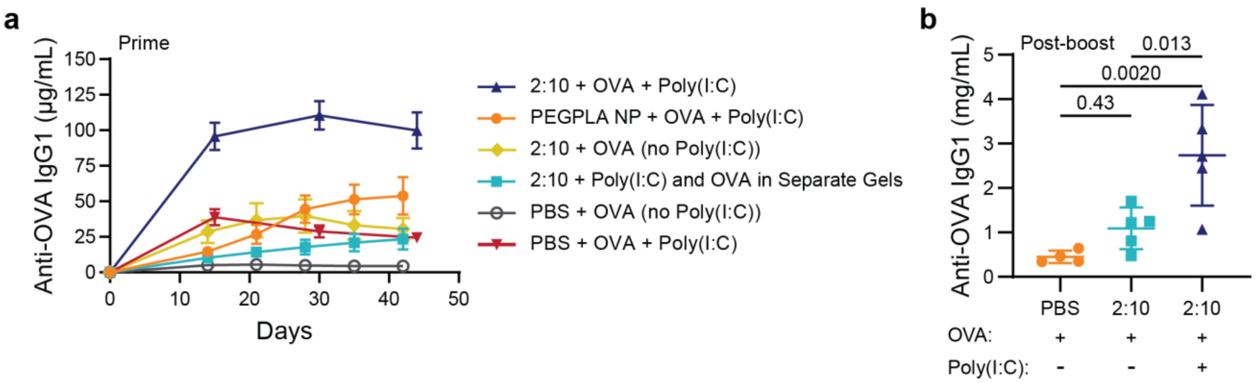

**Figure S5 | Antibody concentrations after prime and boost for various vaccine formulations. a**, Serum anti-OVA IgG1 concentrations from day 0 to day 42 or 45 after prime of vaccines (n = 5 to 19) delivered in different formulations (Error bars, mean ± s.e.m.): (i) OVA and Poly(I:C) in a 2:10 gel, (ii) OVA and Poly(I:C) with PEGPLA NPs, (iii) OVA alone in a 2:10 hydrogel, (iv) OVA and Poly(I:C) in separate 2:10 gels administered on contralateral flanks, (v) OVA alone in PBS, and (vi) OVA and Poly(I:C) in PBS. **b**, Serum anti-OVA IgG1 concentrations 15 days after a day 45 boost (Error bars, mean ± s.d.). All groups were boosted with a bolus vaccine with OVA and Poly(I:C) to assess their responsiveness on the same timeframe. These results show that co-delivery of OVA with the Poly(I:C) adjuvant significantly increases the humoral immune response. Statistical analysis, $*p < 0.05$, $**p < 0.01$ with one-way ANOVA.

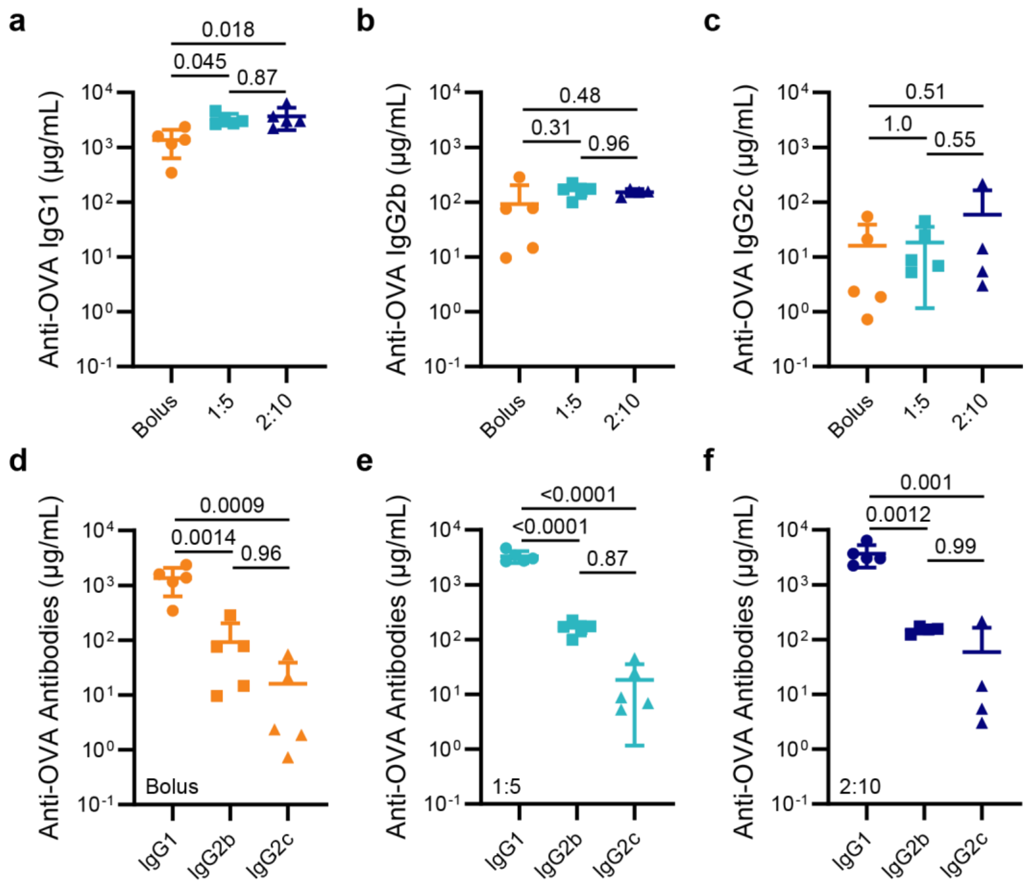

**Figure S6 | Characterization of IgG subclasses in OVA+Poly(I:C) vaccine post-boost.** Serum anti-OVA IgG1 (**a**), IgG2b (**b**), and IgG2c (**c**) concentrations for the bolus, 1:5 gel, and 2:10 gel vaccine groups 15 days after a day 90 bolus boost. Data reorganized for the bolus (**d**), 1:5 gel (**e**), and 2:10 gel (**f**) to show additional statistical comparisons. P values determined by one-way ANOVA with Tukey's test.

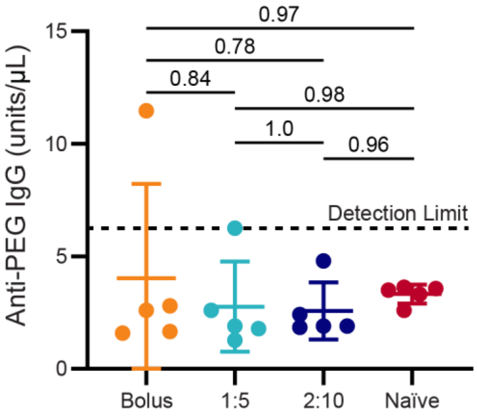

**Figure S7 | Anti-PEG antibody response after OVA+Poly(I:C) vaccine after boost**. Anti-PEG IgG in serum 15 days after bolus boost on day 90 for bolus, 1:5 gel, and 2:10 gel primes along with naïve serum for comparison. All samples were either below or near the detection limit of the assay and no significant differences were found between treatments and naïve serum. P values determined by one-way ANOVA with Tukey’s test.

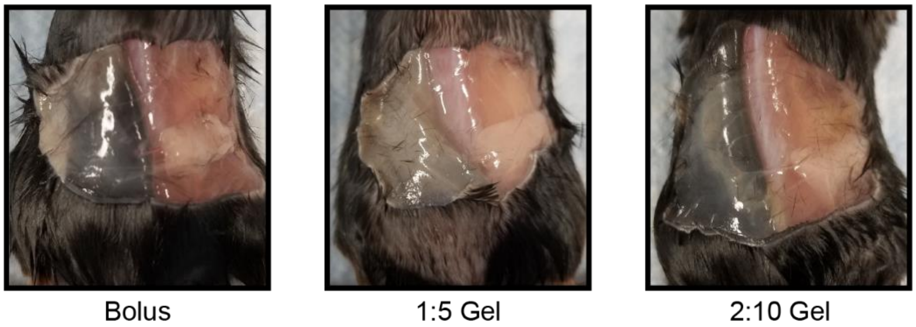

**Figure S8 | Long-term biocompatibility in subcutaneous space**. Images of the subcutaneous space 8 weeks after OVA+Poly(I:C) vaccine administration for the bolus, 1:5 gel, and 2:10 gel treatment groups. Images indicate no noticeable vascularization or fibrotic response differences. The hydrogel materials were completely degraded by this time point.

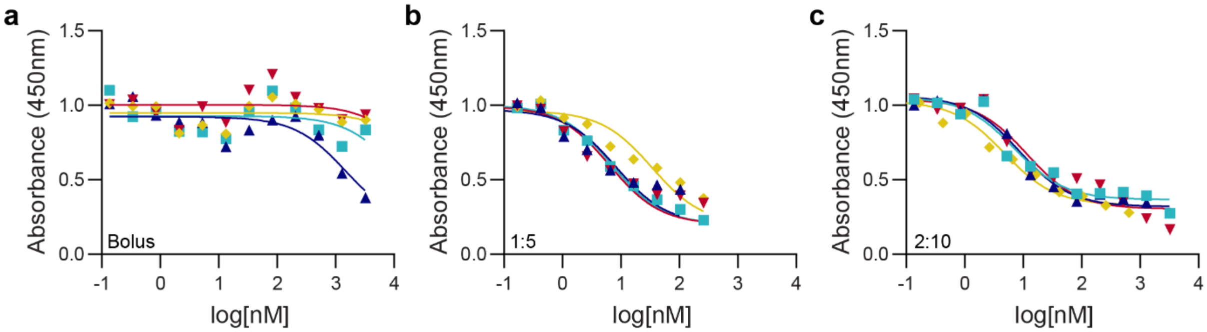

**Figure S9 | Competitive binding assay with serum 15 days after a day 90 boost**. Individual binding curves for all bolus (**a**), 1:5 (**b**), and 2:10 (**c**) samples with the competitive binding fits used to calculate the $K_D$.

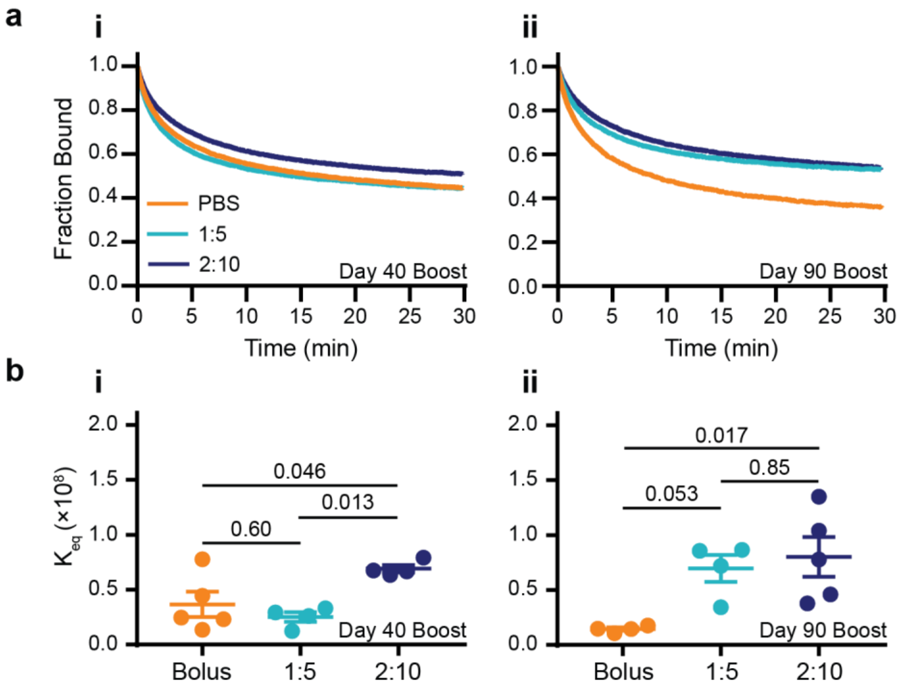

**Figure S10 | Surface plasmon resonance (SPR) affinity analysis of anti-OVA serum antibodies post-boost**. **a**, OVA dissociation from serum antibodies measured by SPR after a day (i) 40 or (ii) 90 boost (n=4). **b**, $K_{eq}$ values for high-affinity antibodies from SPR for a day (i) 40 and (ii) 90 boost using the lowest $k_d$ from a 3-decay fit of the dissociation and the measured $k_a$ (n = 4 to 5). Severe limitations arise in measuring dissociation times with this instrument, which can only collect dissociation data for roughly 30 min (corresponding to $k_d$ values of ~$10^{-4}$ $s^{-1}$, and thus $K_{eq}$ values of ~$10^8$ $M^{-1}$) before baseline drift introduces significant error into the measurement. These data, therefore, serve primarily to verify the trends observed with the more accurate competitive binding assay reported in Figure 3 in the manuscript. See Supplementary methods for more details. P values determined by one-way ANOVA and Tukey's post hoc test.

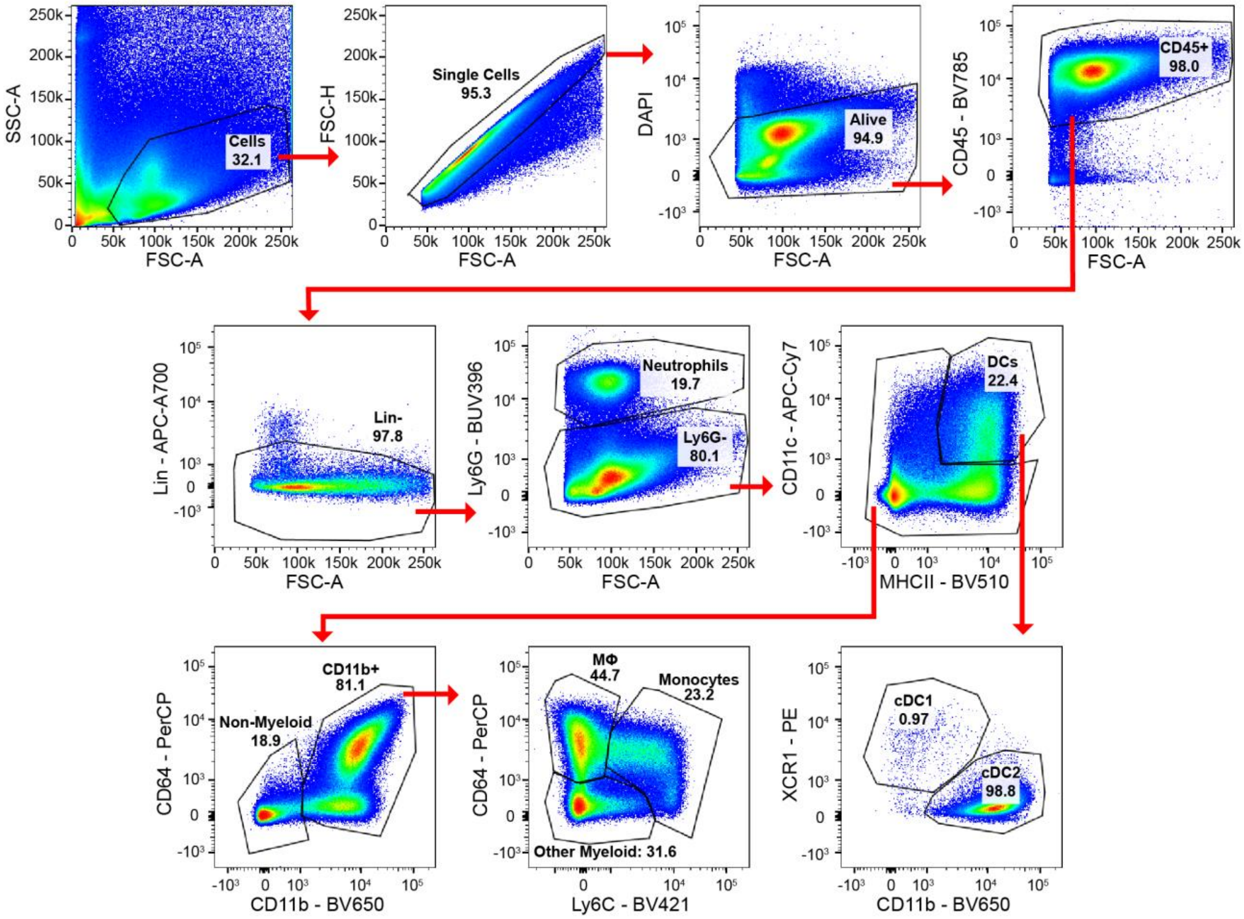

**Figure S11 | Representative gating strategy for gel infiltration analysis.** Neutrophils were defined as CD45+ CD19- CD3- Ly6G+. Dendritic cells were defined as CD45+ CD19- CD3- Ly6G- MHCII+ CD11C+ with cDC1s as XCR1$^{hi}$ CD11b$^{lo}$ and cDC2s as XCR1$^{lo}$ CD11b$^{hi}$. Monocytes were defined as CD45+ CD19- CD3- Ly6G- CD11b+ Ly6C+. Macrophages were defined as CD45+ CD19- CD3- Ly6G- CD11b+ Ly6C- CD64+.

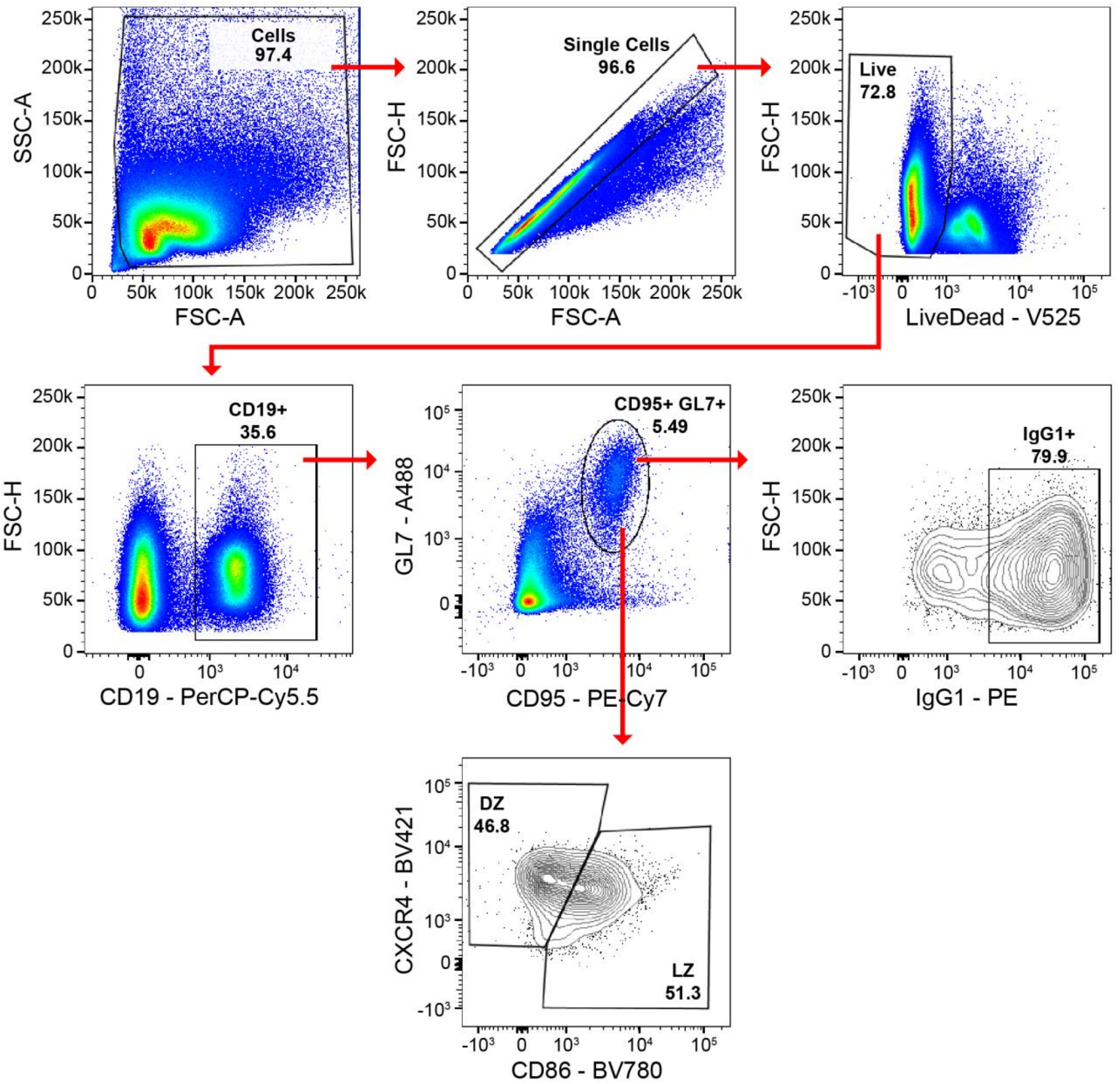

**Figure S12 | Representative gating strategy for GC analysis**. GCBCs were defined as CD19+ CD95+ GL7+.

#### Supplemental Discussion

##### Discussion S1 | Cargo diffusivity in PNP

The diagrams and diffusivity values in SI Fig. 4 show that in PBS the molecular size of a given cargo dictates its diffusivity according to the Stokes-Einstein equation, leading to approximately 10-fold different diffusivities for OVA and Poly(I:C) since they are approximately 10-fold different in size. When entrapped in a covalent PEG gel, the smaller cargo moves freely while the much larger Poly(I:C) is hindered by the mesh of the polymer network, leading to an even larger discrepancy in the diffusivities of the two cargo. In contrast, when encapsulated within the 1:5 and 2:10 PNP gels, both OVA and Poly(I:C) are hindered by the polymer network and their diffusion is therefore limited by the self-diffusivity of the dynamic PNP hydrogel network. Indeed, both cargo exhibit significantly slower rates of diffusion than are observed in PBS because of the obstruction the polymer network poses to their diffusion. When encapsulated in the 2:10 gel in particular, these two biomolecules exhibit matched diffusivities, despite their approximate 10-fold difference in size, as both cargo are completely hindered by the PNP hydrogel matrix. These findings suggest that when a hydrogel matrix is constraining to the diffusivity of entrapped cargo, the hydrogel self-diffusivity can be used as a powerful design principle for precisely engineering desired cargo diffusion properties within dynamically crosslinked supramolecular hydrogels.

##### Discussion S2 | Germinal center response overview

Germinal centers (GCs) are sites within lymphoid organs where mature B cells undergo somatic hypermutation (SHM) leading to higher affinity antibodies. Within the GCs, mature B cells go through cycles of proliferation, mutation, and selection in order to create B cells with the highest affinity B cell receptors (BCRs). The GC is divided into two anatomical compartments: the light zone (LZ) and the dark zone (DZ). In the LZ, GC B cells compete to capture antigen from follicular dendritic cells (FDCs) and present antigen peptides on MHCII[2]. The GC B cells with the highest affinity BCRs have increased antigen presentation to the T follicular helper (Tfh) cells and therefore will receive more positive signaling from Tfh cells. Tfh cell signaling is critical for positive selection of B cells with BCRs which have high affinity for antigen. The B cells that successfully interact with Tfh either mature into plasma cells or return the DZ for another round of proliferation and additional SHM[2,3]. For this reason, prolonged Tfh cell presence in GCs is indicative of prolonged SHM. Similarly, a shift towards LZ B cell markers indicates an increase in the process of affinity selection compared to the processes of expansion and SHM which is necessary to continuing B cell cycling through the GC[4].

#### Supplemental Tables

##### Table S1 | Nanoparticle Characterization (Measured with DLS)

| | Diameter (nm) | Zeta Potential (mV) |
|---|---|---|
| **PEG-PLA NP (n=4)** | 32 ± 4 | -28 ± 7 |

**Table S2 | Cargo and Polymer Diffusivities (Measured with FRAP)**

| Sample | Diffusivity (μm$^2$/s) |
|---|---|
| Poly(I:C) in PBS* | 7.2 |
| OVA in PBS* | 76.6 |
| Poly(I:C) in 1:5 | 1.7 ± 0.3 |
| OVA in 1:5 | 6 ± 1 |
| HPMC-$C_{12}$ in 1:5 | 0.9 ± 0.1 |
| Poly(I:C) in 2:10 | 2.4 ± 0.1 |
| OVA in 2:10 | 3.2 ± 0.6 |
| HPMC-$C_{12}$ in 2:10 | 0.68 ± 0.08 |
| Poly(I:C) in PEG Gel** | 0.4 |
| OVA in PEG Gel** | 66.2 |

*Calculated with Stokes-Einstein (details in methods)

** Calculated with Multiscale Dynamic Model (details in methods)

**Table S3 | Flow cytometry antibody information**

| Antibody (all anti-mouse) | Manufacturer | Clone |
|---|---|---|
| CD19-PerCP-Cy5.5 | BioLegend | 6D5 |
| GL7-A488 | BioLegend | GL7 |
| CD95-PE-Cy7 | BD Biosciences | Jo2 |
| CXCR4-BV421 | BioLegend | L276F12 |
| CD86-BV785 | Prepared in Pulendran Lab | GL1 |
| IgG1-PE | BD Biosciences | A85-1 |
| CD4-BV650 | BioLegend | GK1.5 |
| CXCR5-BV421 | BioLegend | L138D7 |
| PD1-A647 | BioLegend | 29F.1A12 |
| CD45-BV785 | BioLegend | 30-F11 |
| CD3-BV650 | BioLegend | 17A2 |
| CD19-Alexa700 | BD Biosciences | 1D3 |
| CD11c-APC-Cy7 | ebioscience | N418 |
| Ly6G-BUV395 | BD Biosciences | 1A8 |
| CD11b-BV650 | BioLegend | M1/70 |
| MHCII-BV510 | BioLegend | M5/114.15.2 |

#### Supplemental Video

**Video S1 | PNP Hydrogel Injection.** Video of 2:10 PNP hydrogel being injected through a 21-gauge needle. The hydrogel exhibits high viscosity before injection, but dramatically shear-thins and flows through the needle during injection, and rapidly returns to a high viscosity, shape-persistent state once exiting the needle.

#### Supplemental Methods

##### Materials

All chemicals were obtained from Sigma-Millipore, unless specified otherwise.

##### FRAP Analysis

Alexa Fluor 647 conjugated OVA (Thermo Fisher Scientific), rhodamine conjugated Poly(I:C) (Invivogen), and rhodamine conjugated HPMC-$C_{12}$ were used to visualize the diffusion of the cargo and gel. The samples were placed in a sterile 0.18 mm thick glass bottom dish (Ibidi). An Inverted Zeiss LSM 780 Laser Scanning Confocal Microscope (Germany) using a Plan-Apochromat 20X/0.8 M27 objective lens was used for FRAP measurements. To excite the fluorescent Alexa Fluor 647, a 5 mW 633 nm He-Ne laser was employed at 2%, and the emitted fluorescence was detected by Alexa Fluor 647 specific band pass filter (638 – 756 nm). To excite the fluorescent rhodamine, a 20 mW 561 nm diode pumped solid state laser was used at 2%, and the emitted fluorescence was detected by rhodamine specific band pass filter (415 – 638 nm). We used pixel dwell time 2.55 µs which took 390.98 ms to finish every scan. Samples were photobleached with a 405 nm diode laser, plus the 633 nm laser in the case of the Alexa Fluor, and a 514 nm argon laser and the 561 nm laser in the case of the rhodamine. All lasers were set at 100% intensity for the bleaching, with a 20-40 µm diameter for the region of interest (ROI). To avoid any extra noise, the high voltage was limited to be 700 V. Different tests (n = 5) were made for 3 different samples from the same batch at different locations of the sample. For each test, 10 control pre-bleach images were captured at 1 frame/s. A spot was bleached with a pixel dwell time of 177.32 µs. 500 post-bleach frames were recorded at 1 frame/s to form the recovery exponential curve. The pixel size was set to be 1.66 µm. The diffusion coefficient was calculated as[5]:

$$D = \gamma_D(\omega^2/4\tau_{1/2})$$

Where the constant $\gamma_D = \tau_{1/2}/\tau_D$ , with $\tau_{1/2}$ being the half-time of the recovery, $\tau_D$ the characteristic diffusion time, both yielded by the ZEN software, and $\omega$ the radius of the bleached ROI (25 µm). The software used for all FRAP tests was the ZEN lite (Zeiss). All the experiments were conducted in the Stanford University Cell Sciences Imaging Facility (CSIF) at room temperature.

$R_g$ of Poly(I:C) was obtained from gel permeation chromatography (GPC) carried out using a Dionex Ultimate 3000 instrument (including pump, autosampler, and column compartment). Detection consisted of an Optilab TrEX (Wyatt Technology Corporation) refractive index detector operating at 658 nm and a HELEOS II light scattering detector (Wyatt Technology Corporation) operating at 659 nm. The column used was a Superose 6 increase 10/300 GL. The eluent was PBS buffer, 137 mM NaCl, 0.0027 mM KCl, 10 mM Phosphate pH 7.3, at 0.75 mL min$^{-1}$ at room temperature. Analyte samples at 2 mg mL$^{-1}$ were filtered through a PVDF membrane with 0.2 mm pore size prior to injection. $R_g$ was calculated using a dndc of 0.19 mL/g and converted to $R_H$ using the following equation[6]: $R_g/R_H \approx 0.77$.

The diffusivity of cargo in PBS was calculated using Stokes-Einstein Law Equation for diffusion[7] where $k_B$ is Boltzmann's constant, T is temperature in Kelvin, η is solvent viscosity, and R is solute hydrodynamic radius:

$$D = \frac{k_B T}{6\pi\eta R}$$

The diffusivity of cargo in a model covalent PEG gel was calculated using the Multiscale Diffusion Model (MSDM) assuming 25 °C, 5% volume fraction, and 35 nm mesh size[1]. The calculated values are comparable to experiment diffusivities of similar sized cargo found in the literature[8].

**NMR Spectroscopy**

NMR spectra were obtained using an Inova 300 MHz NMR spectrometer with a Varian Inova console using VNMRJ 4.2 A software. Number-average (Mn) and weight-average (Mw) molar mass and dispersity (Đ = Mw/Mn) of polymers were obtained from gel permeation chromatography (GPC) carried out using an Dionex Ultimate 3000 instrument (including pump, autosampler, and column compartment) outfitted with an ERC Refractomax 520 refractometer. The columns were Jordi Resolve DVB 1000 Å, 5 m, 30 cm x 7.8 mm and a Mixed Bed Low, 5m, 30 cm x 7.8 mm, with a Jordi Resolve DVB Guard Column, 1000 Å, 5m, 30 cm x 7.8 mm, 5 cm x 7.8 mm. DMF with 10 mM LiBr was used as eluent at 1 mL min$^{-1}$ at room temperature. Poly (ethylene glycol) were used to calibrate the GPC system. Analyte samples at 2 mg mL$^{-1}$ were filtered through a nylon membrane with 0.2 mm pore size before injection (20 μL). Data was analyzed using Chromeleon GPC/SEC Software.

**Surface Plasmon Resonance**

A BIAcore X100 instrument (GE Healthcare) was used for all Surface Plasmon Resonance (SPR) kinetics measurements. Anti-mouse IgG antibodies (Cat. BR100838, GE Healthcare) were immobilized to a CM5 chip (Cat. BR100012, GE Healthcare) using ammine coupling according to the manufacturer's instructions. Serum samples were injected at a 1:100 dilution in HBS-EP+ buffer (Cat. BR100826, GE Healthcare) for 140 seconds at a flow rate of 10 μL/min to capture the mouse antibodies onto the chip. For association analysis, 5 different concentrations of OVA in HBS-EP+ buffer (978 nM, 733 nM, 489 nM, 244 nM, 162 nM) were flowed for 90 seconds at 30 μL/min followed by 90 seconds of buffer alone. For dissociation analysis, OVA at 489 nM was run for 90 seconds at 30 μL/min followed by 30 minutes of HBS-EP+ to allow for OVA to dissociate from the antibodies.

For each run, we had two steps of reference subtraction. First, naïve mouse serum was run in parallel to the test serum in the reference cell of the chip and subtracted from all test curves. Second, buffer was run above the captured antibodies with the same settings as the OVA solutions to subtract for temporal changes due to fluid flow. To calculate the association rate constant, GraphPad Prism 7.04 (GraphPad Software) was used to fit a one-phase association model to all 5 concentrations of OVA. The observed k values from those fits were plotted against the concentration and fit with a linear equation. The slope from this line was defined as the association rate for that sample. To calculate the dissociation rate constants and account for the polyclonal population of antibodies in the serum, GraphPad Prism 7.04 (GraphPad Software) was used to fit a 3-component exponential decay to the dissociation curves:

$$Y=Y1*\exp(-K1*X)+(Y2*\exp(-K2*X))+((Y3)*\exp(-K3*X))$$

The initial values used for the model were:

$$Y1 = 0.25, Y2 = 0.25, Y3 = 0.5$$

$$K1 = 0.1, K2 = 0.001, K3 = 0.001$$

The component with the highest fraction and lowest dissociation rate was used for comparing the experimental groups.
